# Cycle network model of Prostaglandin H Synthase-1

**DOI:** 10.1101/2020.08.12.246124

**Authors:** Alexey Goltsov, Maciej Swat, Kirill Peskov, Yuri Kosinsky

## Abstract

The kinetic model of Prostaglandin H Synthase-1 (PGHS-1) was developed to investigate its complex network kinetics and non-steroidal anti-inflammatory drugs (NSAIDs) efficacy in different *in vitro* and *in vivo* conditions. To correctly describe the complex mechanism of PGHS-1 catalysis, we developed a microscopic approach to modelling of intricate network dynamics of 35 intraenzyme reactions among 24 intermediate states of the enzyme. The developed model quantitatively describes interconnection between cyclooxygenase and peroxidase enzyme activities; substrate (arachidonic acid, AA) and reducing cosubstrate competitive consumption; enzyme self-inactivation; autocatalytic role of AA; enzyme activation threshold, and synthesis of intermediate PGG_2_ and final PGH_2_ products under wide experimental conditions. In the paper we provided the detailed description of the enzyme catalytic cycle, model calibration based on a series of *in vitro* kinetic data and model validation using experimental data on the regulatory properties of PGHS-1.

The validated model of PGHS-1 with a unified set of kinetic parameters is applicable for *in silico* screening and prediction of the inhibition effects of NSAIDs and their combination on the balance of pro-thrombotic (thromboxane) and anti-thrombotic (prostacyclin) prostaglandin biosynthesis in platelets and endothelial cells expressing PGHS-1.

## 1. Introduction

Prostaglandin H synthase-1 (PGHS-1) also known as Cyclooxygenase-1 (COX-1) catalyses the first stage in the conversion of arachidonic acid (AA) into prostaglandin H_2_ (PGH_2_) being precursor of physiologically active prostanoids [1]. PGHS-1 together with the other isoform of PGHS, PGHS-2 are the main targets of non-steroidal anti-inflammatory drugs (NSAIDs) in treatment of pain, fever, inflammation, and cardiovascular diseases [2]. Despite the fact that this enzyme has been under thorough investigation for several decades, some details of its catalytic mechanism still remain not completely understood [3]. It is generally recognized that PGHS-1 has two distinct catalytic activities: a cyclooxygenase (COX) activity converting AA to an intermediate product, prostaglandin G_2_ (PGG_2_) and a peroxidase (POX) activity which further converts PGG_2_ into a final product PGH_2_. PGHS-1 catalysis represents a complex network of many oxidized/reduced reactions connecting multiple enzyme intermediates and leading to redox transformation of key catalytic components such as AA binding site (COX site) and heme-prosthetic group (POX site). Another not completely understood phenomena in PGHS-1 catalysis is so-called self-inactivation process, including gradual decay of both COX and POX activities during PGHS-1 catalysis [4]. Commonly, two different catalytic mechanisms have been suggested to account for the range of experimental data observed in PGHS-1 catalysis: Branched-Chain (BC) mechanism [5] and alternative Tightly Coupled (TC) mechanism [6], [7]. In the framework of these two mechanisms, several various mathematical models have been proposed to elucidate the details of PGHS-1 catalysis.

According to the BC mechanism of the PGHS-1 catalysis, supported by much experimental data, PGHS-1 catalyses oxygenation of AA to prostaglandin PGH_2_ in the coupled COX and POX cycles (Fig. 1). At the initial stage of the catalysis, PGHS-1 reacts with hydroperoxide, producing oxidized peroxidase intermediates, the ferryloxo-protoporphyrin radical cation, in the POX site which then undergoes reduction by an electron intra-protein transfer from Tyr385. This leads to the formation of the active enzyme intermediate with Tyr385 radical, the oxidants in the COX site. Tyr385 radical abrogates a hydrogen atom from AA and initiates reaction of AA oxygenation to yield PGG_2_ peroxyl radical (PGG_2_*) which is reduced to hydroperoxide prostaglandin G (PGG_2_) accompanied by regeneration of the Tyr385 radical. The transient product PGG_2_ exits the COX site and the further reaction occurs in the POX site, where PGG_2_ is reduced to the final product, endoperoxyalcohol PGH_2_ by the heme prosthetic group, regenerating oxidized peroxidase intermediates in the POX site. The produced PGG_2_ activates autocatalytically cyclooxygenase catalysis in other PGHD-1 enzymes. Each oxidized peroxidase intermediate in the POX site undergoes a one-electron reduction by an exogenous reducing substrate (RC). In contrast, the TC mechanism of PGHS-1 catalysis suggests that the POX and COX activities are tightly coupled in one cycle, and COX cycles cannot function independently on the POX cycle [6], [7].

**Fig. 1.**
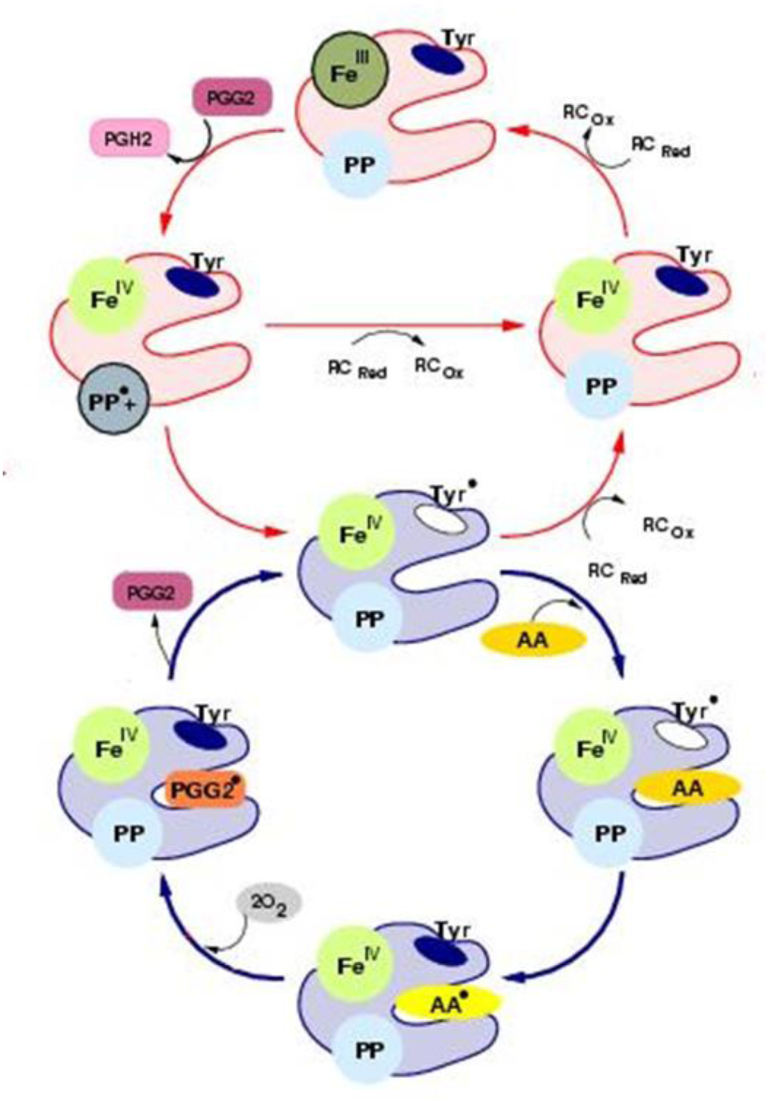
Scheme of the Branched Chain mechanism of PGHS -1 catalysis [5]. Detailed description of all the enzyme states is given in Table 1.

The most detailed mathematical models of PGHS-1 have been developed in a series of papers [5], [8], [9], [10]. These kinetics models were based on the assumptions of BC mechanism of PGHS-1 function, and gradually evolved starting from eight reactions in an initial model [8] to 22 reactions in the latest version [10]. An increase in the complexity of the models reflected a growing insight into the mechanism of catalysis and molecular structure of PGHS-1. An extension of the models included an introduction of additional enzyme intermediates and reactions into these proposed catalytic schemes and more detailed consideration of enzyme self-inactivation. Each step in a gradual refinement of the model and an increase in its complexity were supported by the new experimental data as it became available. The developed models allowed satisfactorily describing key features of PGHS-1 catalysis, i.e. interaction of POX and COX activities, self-inactivation and autocatalytic effect, as well as estimating some kinetic constants of the PGHS-1 catalytic reactions [11].

Abundance of accumulated experimental data on the PGHS-1 kinetics stimulated development of various kinetic models of PGHS-1. Several PGHS-1 models have been developed with a view to describe individual datasets and explain certain peculiarities of PGHS-1 functioning. For example, the kinetic models of PGHS-1 developed by Vrzheshch et al. [12] allowed qualitatively describing sets of experimental data on AA, O_2_ and RC consumption under specific *in vitro* conditions. The kinetic models of two isoforms, PGHS-1 and PGHS-2 were developed to explain the differences in their kinetics and activation thresholds [13].

None of the existing mathematical models of PGHS-1 can provide quantitative description of all key regulatory effects within a unified set of kinetic parameters. In the other words, different models and sets of parameters should be used to reproduce accurately a wide spectrum of experimental data on PGHS-1 kinetics and predict its function *in vivo* in health and pathological conditions. It complicated significantly the use of the existing models of PGHS-1 for *in silico* study of inhibition effect of NSAIDs in different *in vitro* and *in vivo* conditions, and for their application in the development of Quantitative Systems Pharmacology (QSP) models, where prostaglandins pathway is one of the drug targets.

In spite of great progress in understanding of PGHS-1 functioning, the problem of quantitative description of the main experimental facts on its reaction mechanism as well as the reproduction of experimental data obtained in different *in vitro* and *in vivo* conditions in the framework of a unified kinetic model is still remaining. The new experimental data appears and adds new knowledge on the enzyme structures and functions which can play a role in prostaglandin biosynthesis [14].

Basing on the ideas of BC mechanism [5] and their realisation in the kinetic models [8]– [10], in this paper we developed the microscopic kinetic approach to model the complex network dynamics of intra-enzymatic reactions among multiple catalytically active intermediate states of the enzyme and reproduce the detailed kinetic mechanisms of the PGHS-1 function. Note that this model has been previously applied to the *in silico* description of NSAID action alone and in combination and showed satisfactory predictive results [15], [16]. In particular, the model was applied to the investigation of dose dependence of PGHS-1 for aspirin and selective NSAID, celecoxib, on concentrations of arachidonic acid, reducing cosubstate and hydrogen peroxide which were reported to differ dramatically in purified and cellular assays as well as in *in vivo* conditions in different cells and organs [17]. The model also was applied to the analysis of decreasing aspirin efficacy at its administration in combination with selective NSAIDs [15], [16]. But many aspects of the model and features of the interaction of NSAIDs and PGHS-1 have remained beyond the scope of these publications. In this paper we give a full description of the model assumptions, method of its parametrisation, validation, and model application to the analysis of experimental data on regulatory properties of PGHS-1. One of the main aims of this work is the construction of reliable and validated kinetic model of PGHS-1, which would incorporate current theoretical conceptions on the PGHS-1 function and would be capable to reproduce quantitatively much experimental data on prostaglandin biosynthesis in the framework of a unified set of kinetic parameters.

## 2. Results

### 2.1. Development of the cycle network model of PGHS-1

The derivation of the model of PGHS-1 catalysis represents a special task, as both POX and COX activities are known to be located in different enzyme domains within the same protein complex, and these activities can proceed independently of each other [18]. Another important feature of PGHS-1 catalysis is that it involves intramolecular electron transfer and formation of multiple redox and binding states of the catalytic domain. The start point of our modelling was the BC mechanism [5], [9] schematically represented in Fig.1. The full catalytic cycle of PGHS-1 includes two distinct catalytic activities: a cyclooxygenase (COX) activity converting arachidonic acid (AA) to an intermediate product, prostaglandin G2 (PGG_2_), and a peroxidase (POX) activity, which further converts PGG_2_ into final product PGH_2_. Following the approach to the PGHS-1 modelling [9], we assumed that PGHS-1 catalysis could be presented as a network of microscopic reactions, each of them resulting in a corresponding change in the enzyme catalytic state. Under catalytic state we understand the state of PGHS-1 catalytic domain, which in its turn is defined by a unique combination of the states of its key elementary catalytic constituents, located in the COX and POX active sites. In our model we considered the following key catalytically active components: fatty acid binding site (COX site), heme prosthetic group (POX site) and Tyr385 residue in the COX site.

We defined the COX site microstates through the presence/absence of AA in the hydrophobic channel of COX region. The POX state was determined by the redox states of heme iron and protoporphyrin group. Tyr385 residue was assumed to exist in either ground or radical state. In addition, we considered states of fully inactive enzyme, FIE [19]. The detailed description of all-possible states for each catalytic component, considered in our model, is given in Table 1.

**Table 1.** Microstates of PGHS-1 considered in the model.

| **n** | **Catalytic components** | **N** | **Notation of the states** | **Description of the states** |
| --- | --- | --- | --- | --- |
| 1 | Heme prosthetic group (POX site) | 1 | Fe(III), PP | Protoporphyrin group (PP) in ground state; |
|  |  | 2 | Fe(IV), PP*+ | Oxiferryl state of Fe (Fe(IV)=O) and radical cation state of PP group; |
|  |  | 3 | Fe(IV), PP | Oxiferryl state of Fe (Fe(IV)=O) and reduced PP group |
| 2 | Tyrosine 385 | 1 | Tyr | Tyrosine residue |
|  |  | 2 | Tyr* | Tyrosyl radical |
| 3 | AA binding site (COX-site) | 1 | empty | No ligands in the COX-site |
|  |  | 2 | AA | Arachidonic acid in the COX site |
|  |  | 3 | AA* | Arachidonyl radical in the COX site |
|  |  | 4 | PGG<sub>2</sub>* | Prostaglandin G<sub>2</sub> radical in the COX site |
|  |  | 5 | FIE | Fully Inactive Enzyme |

Fig. 2 shows a kinetic scheme for the developed catalytic cycle of PGHS-1. The scheme is mainly based on the assumptions of the BC mechanism [5] and its further extension [9]. Additionally, we also included in the model some reactions corresponding to an early version of the TC mechanism of PGHS-1 catalysis [6]. Thus, our scheme represents a generalisation of two previously discussed catalytic mechanisms, and the results obtained in the framework of the model allowed us to draw the conclusions on the role of each mechanism in PGHS-1 catalysis.

**Fig. 2.**
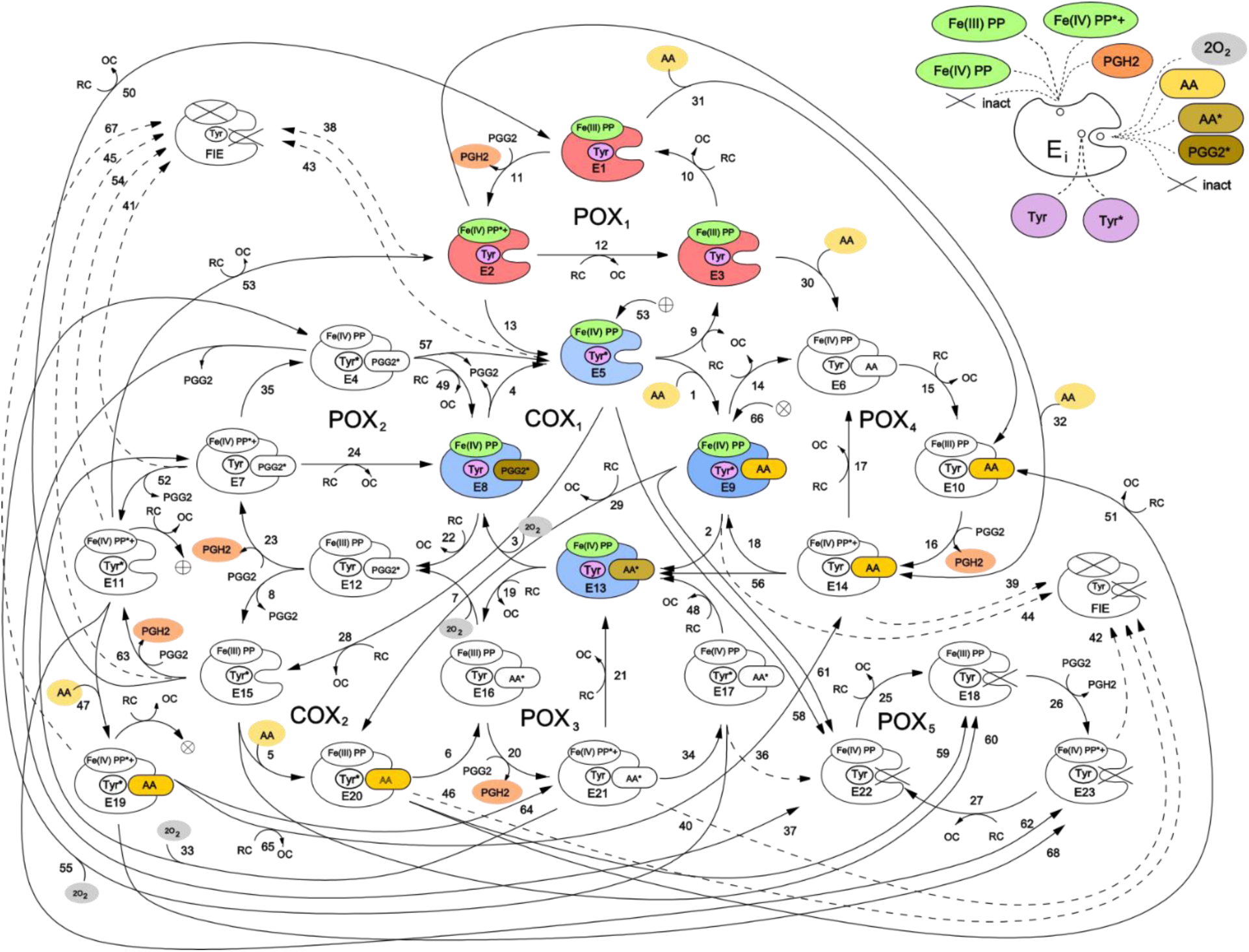
Cycle network model of PGHS-1. Detailed description of all the enzyme states is given in Table 1. Enzyme states marked by colour correspond to the Branched Chain mechanism [5].

We have presented the full catalytic cycle of PGHS-1 as a network of eight interlinked cycles, five of them (POX_i_) corresponding to the POX activity, the other three (COX_j_) – to the COX activity of the enzyme. In general, the full catalytic cycle of PGHS-1 can be considered as a complex dynamic network of elementary reactions which describes the totality of possible transitions between multiple states of the PGHS-1 catalytic domain.

Two interconnected cycles, COX_1_ (reactions 1-4) and POX_1_ (reactions 9-13) correspond to the BC mechanism. According to this mechanism, PGHS-1 catalysis is initiated in the POX site, and its first step is the reaction of the resting enzyme state [Fe(III), PP] (E_1_) with PGG_2_ producing PGH_2_ and leaving the heme prosthetic group in the state [Fe(IV), PP*+] (E_2_). The enzyme state containing heme with two oxidizing equivalents – one on the iron and another one on the porphyrin radical cation – is referred to as Intermediate 1 [5]. In our model, we described this stage of POX catalysis by bimolecular irreversible reaction (reaction 11) of the resting state with PGG_2_, resulting in production of PGH_2_ and leaving the enzyme in the E_2_ with heme in [Fe(IV), PP*+] state. E_2_ state can be converted back to the resting state E_1_ by 2 consecutive reactions with RC by followed release of oxidized cosubstrate, OC (reactions 12 and 10). Alternatively, E_2_ state can undergo one-electron internal reduction via intramolecular electron transfer from Tyr385 residue in the COX site to protoporphyrin (reaction 13) resulting in the formation of the E_5_ state [Fe(IV), PP, Tyr^*^] containing Tyr385 radical (Tyr385*) and a ferryl heme. This state is referred to as Intermediate II [5]. E_5_ state can also complete POX cycle and return to the resting state E_1_ via reactions 9 and 10 with RC through the formation of [Fe(IV), PP] referred as Intermediate II [5]. According to the BC mechanism, the presence of Tyr385 radical is ultimately required for AA oxidation in the COX site. Therefore, the intermediate state E_5_ can potentially serve as a starting point for the further cascade of COX reactions.

The majority of previously known models included COX activity as a single Michaelis-Menten type reaction without considering its mechanism in any detail [10], [13]. In contrast, in our model we describe COX activity by four consecutive reactions (reactions 1-4 in COX_1_ cycle) corresponding to the transitions of the enzyme through three intermediate enzyme-substrate complexes (E_9_, E_13_ and E_8_). Note that using of Michaelis-Menten approximation to describe the individual reactions involved in a reaction network is not strongly valid.

Reaction 1 describes reversible binding of AA to the enzyme state, E_5_ resulting in the formation of the state [Fe(IV), PP, Tyr^*^, AA] (E_9_), containing AA in the COX site. Reaction 2 corresponds to the abstraction of a hydrogen atom from AA by Tyr385 radical and formation of the E_13_ state containing AA radical in the COX site, [Fe(IV), PP, Tyr, AA*]. Further, AA^*^ reacts with two oxygen molecules and the intermediate product PGG_2_ radical (PGG_2_*) is produced (reaction 3). We assumed that binding of the first and second oxygen molecules is a rapid process, during which the redox state of the heme remains unchanged [8]. The final step of the COX_1_ cycle in our model is irreversible reaction 4, which describes the process of regeneration of Intermediate II with Tyr385 radical (E_5_), accompanied by the release of the intermediate product PGG_2_ from the COX site.

Taking into account that POX activity of PGHS-1 is observed to proceed independently from COX catalysis [18], we considered the possibility of several additional POX and COX catalytic cycles, originating from intermediate states of the POX_1_ and COX_1_ cycles. For example, in the COX_1_ cycle, heme prosthetic group remains in the state [Fe(IV), PP] with one oxidising equivalent on iron, and therefore each of the enzyme states E_5_, E_8_, E_9_, and E_13_ can potentially participate in redox reactions with RC (reactions 19, 22, 28, and 29), producing catalytic states E_12_, E_15_, E_16_, and E_20_. The later states can be considered as intermediates of an additional COX_2_ cycle. Note that similar COX cycle was included in the model developed by Kulmacz et al. [18]. The COX_2_ cycle differs from the COX_1_ cycle only by the state of heme prosthetic group in the enzyme intermediates. Similarly, each intermediate of POX_1_ cycle has an empty COX binding site which allows for binding of AA (reactions 30, 31 and 32) and production of the states E_6_, E_10_ and E_14_, respectively. These states can participate in the POX_2_ cycle which differs from the POX_1_ cycle by presence of AA in the COX binding site of its enzyme intermediates. Note, that binding of AA to the resting state (E_1_), containing Tyr385 in the ground state, was considered in the PGHS-1 model for the first time, as all previously known models assumed, that AA binds solely to the enzyme containing Tyr385 radical state (Intermediate II, E_5_). This allowed us to analyse how the redox state of key catalytic components can influence the enzyme binding to AA.

The third COX_3_ cycle though not explicitly shown in the scheme shown in Fig. 2, can be defined as a cycle formed by the enzyme states E_11_, E_19_, E_21_, and E_7_, involving in reactions 47, 64, 33, and 53, respectively. The reactions in the COX_3_ cycle are initiated in the result of AA binding to the enzyme being in two-radical state [Fe(IV), PP*+, Tyr^*^] (state E_11_). Note, that similar COX cycle was considered in the extended BC mechanism model [10], [18], but its contribution to PGHS-1 functioning was not indicated.

Using similar approach, we introduced into our model the POX_3_ and POX_4_ cycles, intermediates of which differ from POX_1_ intermediates by the occupancy of the COX binding site – it contains AA radical in the POX_3_ cycle and PGG_2_ radical in the POX_4_ cycle. The POX_5_ cycle was included to account for remaining POX activity, when the COX site is inactivated (see more detailed description below).

In our model, we also included an alternative route for initiation of COX reactions, which was proposed in an early version of the TC mechanism of PGHS-1 function [5] - a direct hydrogen atom abstraction from AA by protoporphyrin radical cation PP*+ and generation of AA radical. In our scheme this mechanism is described by the reaction of AA binding to the initial state E_1_ resulting in the production of enzyme-substrate complex E_14_, [Fe(III), PP, Tyr, AA], which further converts into intermediate, [Fe(IV), PP, Tyr, AA*] (E_13_), containing arachidonyl radical, AA* (reaction 56). Consideration of two alternative mechanisms of AA radical formation in our model allowed us to analyse their possible contributions to overall COX activity.

### 2.2. Modelling of PGHS-1 self-inactivation

PGHS-1 activity undergoes irreversible self-inactivation (SI) during catalysis [19]. The exact mechanism of this process remains not completely understood [3]. However, there is a consensus that SI most likely originates from the enzyme damage caused by oxidative reactions, coming from certain radical intermediates generated in PGHS-1 [20]. Potentially, any enzyme intermediate containing oxidising equivalents or radicals can cause the enzyme damage. One of the key questions discussed in the literature is which intermediate states can serve as the origin for POX and COX inactivation.

Based on the experimental data we considered several possible mechanisms for enzyme SI in our model. These mechanisms included separate inactivation of the COX and POX activities, originating from enzyme intermediates with different redox state of heme group and containing one or two radicals in the catalytic domains [20]. All the SI processes accounted for in our model can be conditionally classified into the two following groups.

First, inactivation of POX activity resulting in fully inactivation enzyme (FIE state in Fig. 2) was taken into consideration. According to the BC mechanism, SI of POX activity in leads to the subsequent loss of COX activity, as POX reactions are required to regenerate Tyr385 radical state. In our model, this SI mechanism is described by reactions 43-46 and 54. Due to simplicity reasons, we did not distinguish two stages in this process and approximated it by the following one-stage irreversible first order reactions:

a. Reactions 38-42, 53, and 67 originate from Intermediate I and lead to POX and further COX inactivation due to radical state of heme prosthetic group ([Fe(IV), PP*+]) [21], [22]. In our model this SI process can be initiated from seven catalytic states containing radicals such as E_2_, E_7_, E_11_, E_14_, E_19_, E_21_, and E_23_;
b. Reactions 43-46 originate from Intermediate II (states E_5_, E_9_, E_15_, and E_20_) and lead to POX and further COX inactivation due to the presence of oxidising equivalent Fe(IV) in heme.

Second, we considered in the model direct inactivation of COX activity with remaining POX activity which can be caused by the presence of radical states in the COX site [19]. In our model, the remaining POX activity is described by the POX_5_ cycle (reactions 25-27). Similar POX cycle was considered in the model developed by Kulmacz et al. [18]. According to this hypothesis, we assumed that COX inactivation can originate from the enzyme intermediates containing either one or two radicals in the COX catalytic site. Reactions 58-62, and 68 originate from the states containing one Tyr385 radical in the COX site (E_5_, E_9_, E_11_, E_15_, E_19_, and E_20_), reactions 36 and 37 - from the states with two radicals in the COX site, [Fe(IV),PP, Tyr*, AA*] (E_17_) and [Fe(IV),PP, Tyr*, PGG2*] (E_4_). It should be noted that remaining POX activity can also further be inactivated due to damage caused by presence of radical state in heme (reaction 42).

### 2.3. Identification of a set of kinetic parameters of PGHS-1

As the result of joint fitting of three series of the experimental data (see Methods), a unified set of kinetic parameters was determined. The results of fitting are shown in Fig. 3 and presented in Table 2. For the comparison of the obtained kinetic constants of PGHS-1 with the results of other authors, we also included in Table 2 experimental and theoretical estimations of the corresponding rate constants available from literature.

**Table 2.** Kinetic parameters of the PGHS-1 model.

| Reactions | Kinetic constants | Values | Experimental or modelling data |
| --- | --- | --- | --- |
| *COX cycle reactions* |  |  |  |
| $E_5 + AA$ | $k_1$ | $690 \mu M^{-1} s^{-1}$ | |
| | $K_{d,1}$ | $0.1 \mu M$ | |
| $AA \rightarrow AA^*$ | $k_2$ | $2690 s^{-1}$ | |
| $AA^* + 2O_2$ | $k_3$ | $173 \mu M^{-2} s^{-1}$ | |
| $PGG_2^* + Tyr$ | $k_4$ | $13 s^{-1}$ | |
| *POX reactions with RC (phenol) and PGG<sub>2</sub>* |  |  |  |
| $Tyr^* + RC$ | $k_5$ | $0.18 \mu M^{-1} s^{-1}$ | $0.001-0.1 \mu M^{-1} s^{-1}$ [4] |
| $Fe(IV)PP + RC$ | $k_6$ | $0.58 \mu M^{-1} s^{-1}$ | $0.53 \mu M^{-1} s^{-1}$ (4 °C) [24] |
| $E + PGG_2$ | $k_7$ | $7.1 \mu M^{-1} s^{-1}$ | 14 (1 °C); 35 (20 °C) [5];<br>21 (25 °C) $\mu M^{-1} s^{-1}$ [22] |
| $Fe(IV)PP^* + RC$ | $k_8$ | $0.001 \mu M^{-1} s^{-1}$ | |
| *Fe(IV)PP\* + Tyr → Fe(IV)PP + Tyr\** |  |  |  |
| | $k_9$ | $310 s^{-1}$ | $300 s^{-1}$ [22]; $320 s^{-1}$ [4] |
| *Reactions in POX<sub>2</sub> cycle* |  |  |  |
| $Tyr^* + RC$ | $k_{10}$ | $0.1 \mu M^{-1} s^{-1}$ | |
| $Fe(IV)PP^* + Tyr$ | $k_{11}$ | $1.1 s^{-1}$ | |
| *AA binding to the resting enzyme, E<sub>1</sub>* |  |  |  |
| $E_1 + AA$ | $k_{12}$ | $690 \mu M^{-1} s^{-1}$ | |
| | $K_{d,12}$ | $0.6 \mu M$ | |
| *Fe(IV)PP\* + AA → Fe(IV)PP + AA\** |  |  |  |
| | $k_{13}$ | $18.8 s^{-1}$ | |
| *Self-inactivation reactions* |  |  |  |
| $COX_{inact}$ | $k_{in}$ | $0.72 s^{-1}$ | $0.1 s^{-1}$ [4] |
| $POX_{inact}$ | $k_{in1}$ | $0.6 s^{-1}$ | $0.2 - 0.5 s^{-1}$ (24 °C) [4] |
| $COX_{inact}$ | $k_{in2}$ | $2.5 s^{-1}$ | $0.22 s^{-1}$ [4] |
| *POX reactions with adrenaline and H<sub>2</sub>O<sub>2</sub>* |  |  |  |
| | $k_5$ | $0.06 \mu M^{-1} s^{-1}$ | |
| | $k_6$ | $0.001 \mu M^{-1} s^{-1}$ | |
| | $k_7$ | $0.001 \mu M^{-1} s^{-1}$ | $0.44 \mu M^{-1} s^{-1}$ (6.7 °C) [6],<br>$0.01 \mu M^{-1} s^{-1}$ (1 °C), $0.028$ (20 °C) [5] |
| | $k_8$ | $0.001 \mu M^{-1} s^{-1}$ | |

**Fig. 3.**
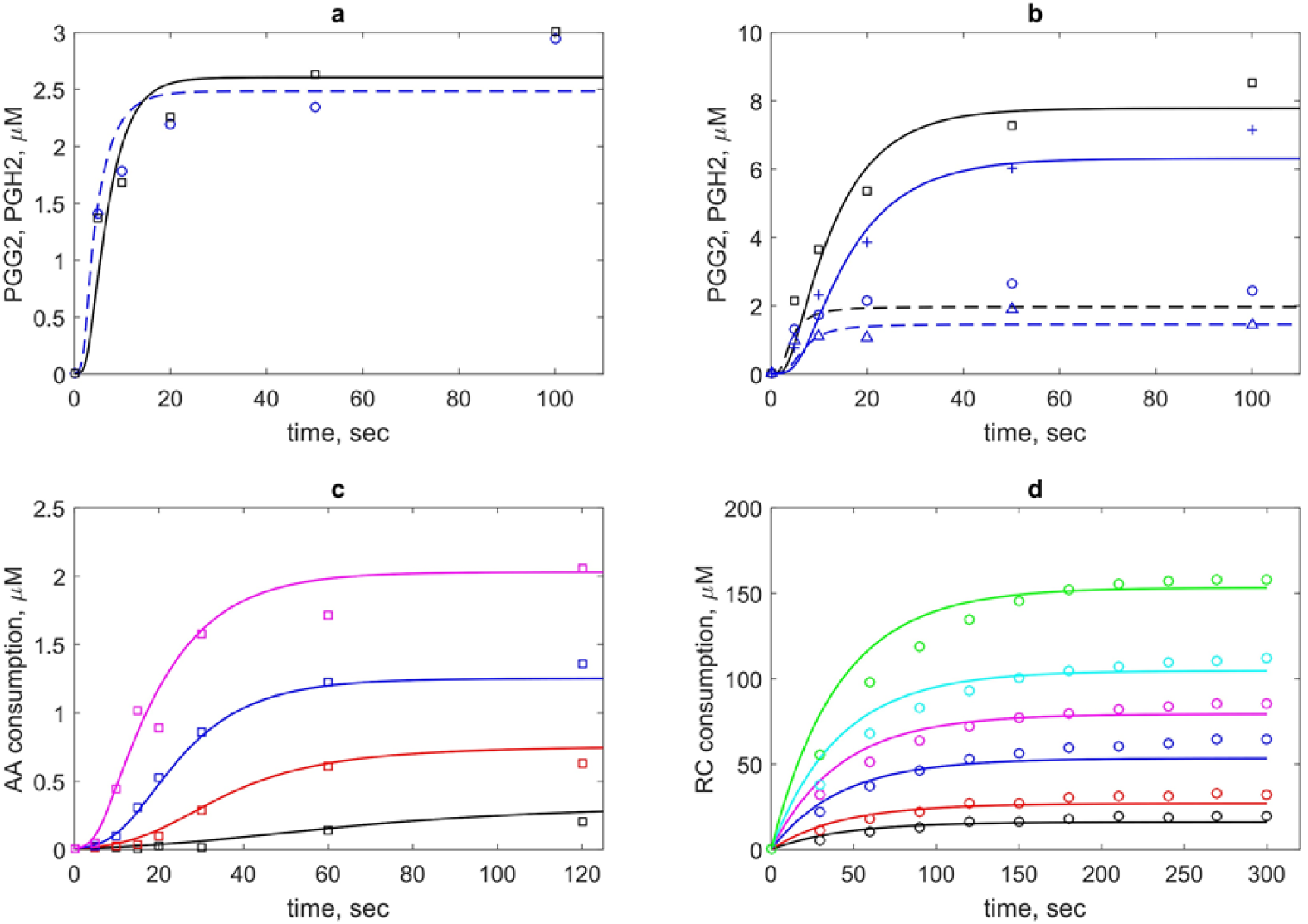
The results of the theoretical description (lines) of experimental data on PGHS-1 kinetics (points). (a) Kinetics of PGG_2_ (dash blue line) and PGH_2_ (black solid line) at 100 μM phenol. (b) Kinetics of PGG_2_ (dash lines) and PGH_2_ (solid lines) at 1000 μM (blue lines) and 5000 μM (black lines). Theoretical and experimental data [8] (a and b) obtained at 80 μM and 35 nM PGHS-1; (c) Theoretical and experimental [11] kinetics of AA consumption at 0.5 μM (black line), 1 μM (red line), 2 μM (blue line), and 20 μM (magenta line) PGHS-1 and 1000 μM phenol. (d|) Accumulation kinetics of oxidised RC (adrenochrome) at 0.32 μM (black line), 0.54 μM (red line), 1.08 μM (blue line), 1.62 μM (pink line), 2.16 μM (cyan line), and 3.22 μM (green line) adrenaline (RC) and 1050 μM H_2_O_2_. Point – experimental data [12].

In the calibration procedure, we used two experimental datasets on PGG_2_ and PGH_2_ production by PGHS-1 treated by AA and phenol as RC [11] (Fig. 3a-3c). As a result of model fitting against the experimental data, the values of the dissociation constants of AA binding to the COX-site, *K*_*d,1*_ (reaction 1 in the COX_1_ cycle) and *K*_*d,12*_ (reaction 12 in the COX_2_ cycle) were estimated (Table 2). The only experimental estimate available for the stage of AA binding to PGHS-1 is Michaelis constant *K*_*m*_ ranged from 3 μM to 5 μM [1]. However, Michaelis constant is an apparent value and generally depends on the concentration of the second PGHS-1 substrate, RC (see discussion below), and therefore cannot be directly compared with *K*_*d,1*_.

One of the key kinetic parameters of the PGHS-1 catalysis is the reaction rate of intramolecular electron transport [23] from Tyr385 to the heme group leading to the formation of Tyr385 radical. The obtained value for constant *k*_*9*_ equal to 310 s^-1^ is in close agreement with theoretical estimates 300 s^-1^ [22] and 310 s^-1^ [4].

Satisfactory quality of the fitting was achieved at the same values of kinetic parameters of COX activity for similar stages in all the three COX_i_ cycles considered in the model. Note that the COX_1_, COX_2_ and COX_3_ cycles differ only by the state of the heme prosthetic group in the POX site (see discussion in section 2.1). The independence of kinetic constants of the COX reactions on the state of the heme group can indicate the absence of cooperativity between the POX and COX sites.

As the result of fitting, we estimated the rate constants for the different mechanisms of PGHS-1 self-inactivation (Table 2). As discussed in section 2.2, we considered three possible mechanisms of SI in the model. For SI originating from Intermediate I ([Fe(IV),PP*+]), we obtained the value of the rate constant *k*_*in1*_=0.6 s^-1^ which is in good agreement with experimental data [4] (Table 2). For the reactions of COX inactivation (reactions 36, 37, 58-62, and 68) with the remaining POX activity (POX_5_ cycle) we obtained rate constant *k*_*in*_=0.72 s^-1^ (Table 2).

A small value of reaction rate *k*_*8*_ of the reduction of protoporphyrin group radical state (Intermediate I, [Fe(IV), PP*+], E_2_) by RC was obtained to be about zero *k*_*8*_ (0.001 μM^-1^ s^-1^) that means that the Intermediate I is rather reduced by intramolecular process of electron abstraction from Tyr385 with radical formation than by exogenous RC.

To estimate uncertainty of the parameters obtained in the fitting procedure of the model against experimental data we carried out global sensitivity analysis (GSA) of the model (see Methods). GSA revealed that the calibrated model showed uniformed sensitivity to the most kinetic parameters (Fig. 4). The model is weakly sensitive to reaction rate *k*_*12*_ of AA binding to the states in the POX_2_ cycle (E_1_, E_2_ and E_3_), while it is sensitive to the dissociation constant *K*_*d,12*_ of the same reactions. The model also showed weak sensitivity to parameter *k*_*in2*_ which corresponds to one of the SI reactions of the enzyme, but it is highly sensitive to the parameters *k*_*in1*_ and *k*_*in*_ characterising the other SI reactions considered in the model.

**Fig. 4.**
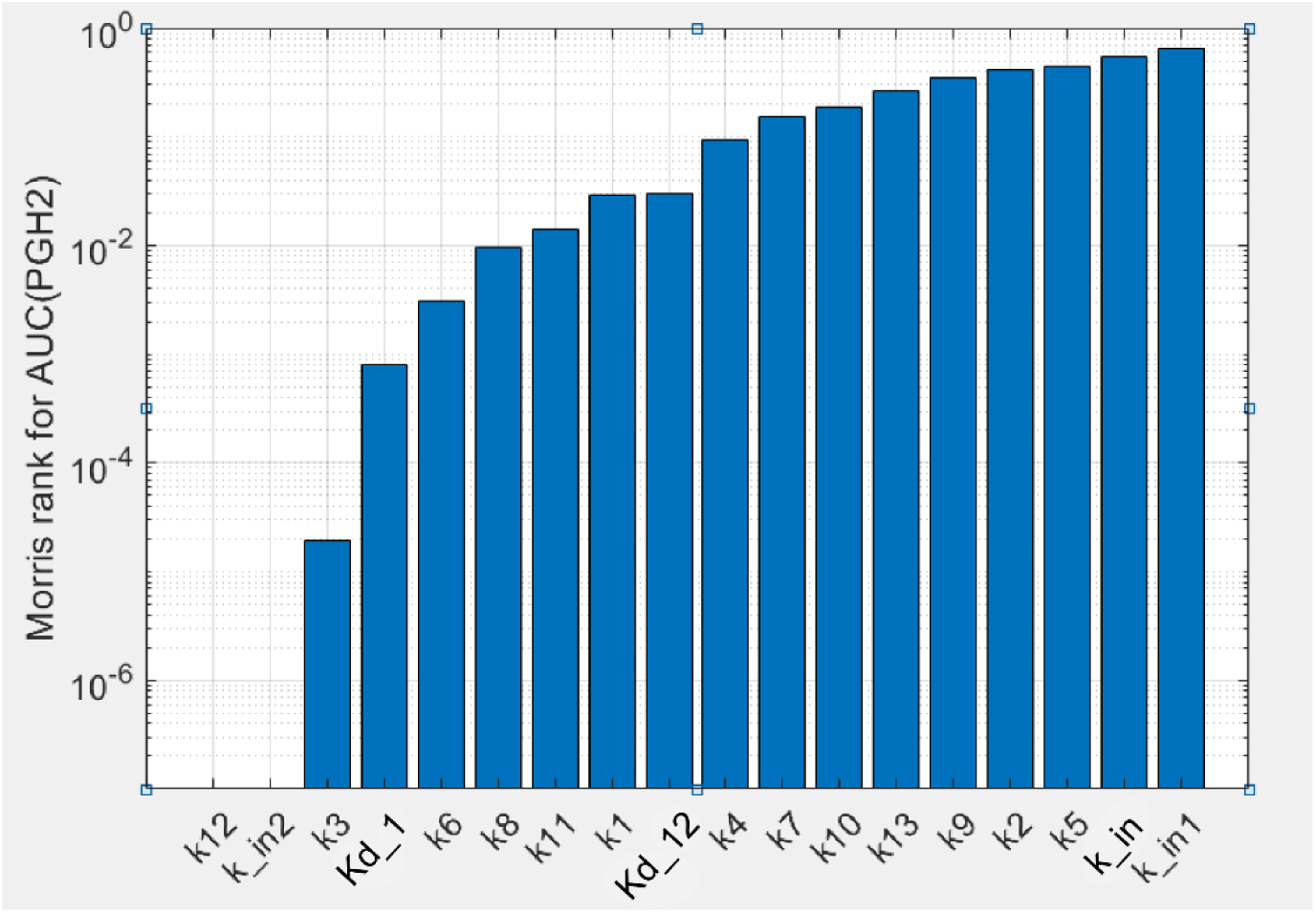
Results of the global sensitivity analysis of the PGHS-1 model. Sensitivity of PGH_2_ production to the kinetic parameters calculated as Morris rank [25] for area under the curve (AUC) of the PGH_2_ time series.

The model also satisfactory described the third experimental dataset on a pure POX activity of PGHS-1 treated by H_2_O_2_ and adrenalin as RC [12] (Fig. 3d). As the result of fitting, another set of the kinetic parameters (*k*_*5*_, *k*_*6*_ and *k*_*8*_) of the POX cycle was obtained which differs from that identified for the POX reactions in PGHS-1 treated by AA and phenol as RC (Table 2). This result is consistent with experimental data on the pure POX activity of PGHS-1 in the presence of different peroxides and cosubstrates [5], [6].

As seen from obtained agreement between theoretical and experimental data (Fig. 3), the model with the unified set of kinetic parameters satisfactorily described the various experimental data such as kinetics of intermediate PGG_2_ and final PGH_2_ products, as well as consumption of substrate, AA and cosubstrate, RC. The model with the obtained set of parameters given in Table 2 was further validated and applied to analysis of regulatory mechanisms of PGHS-1 catalysis.

### 2.4. Contributions of the different enzyme intermediates and intra-enzymatic reactions to PGHS-1 catalysis

The catalytic cycle of PGHS-1 (Fig. 2) represents a complex cycle network of intermolecular states with intricate distribution of internal fluxes between them. To realise their contributions to the PGHS-1 catalysis, we carried out a comparative analysis of the population distribution of intermediate enzyme states and fluxes. This allowed us to estimate relative impact of different elementary reactions and enzyme intermediates to the overall PGHS-1 function. We determined preferred ways of AA binding to the enzyme, identified key reactions producing intermediate PGG_2_ and final product PGH_2_, estimated the contribution of the different POX and COX cycles as well as compared the role of different mechanisms of enzyme SI.

As mentioned above, in the model, we considered two possible ways of initiation of the COX reaction, namely two alternative ways of formation of the enzyme-substrate complex containing both Tyr385 radical and AA. The first way corresponds to a generally accepted mechanism of the COX reaction initiation in the BC mechanism, when formation of Intermediate II state [Fe(IV), PP, Tyr*] (E_5_) precedes the binding of AA (reaction 1). The second, alternative mechanism is the binding of AA to the COX site (reactions 30-32) which can be followed by formation Tyr385 radical state, E_9_ through reaction 18 (Fig. 2).

The performed flux analysis showed that AA binds to the Intermediate II (E_5_) state much more effectively than to the resting state (E_1_), Intermediate I (E_2_) and E_3_ state. Maximum flux *V*_*1*_*(t)* exceeds maximum values of *V*_*30*_*(t), V*_*31*_*(t)* and *V*_*32*_*(t)* by an order of magnitude (see comparison of *V*_*1*_*(t)* and *V*_*31*_*(t)* in Fig. 5a. Note, that the reaction rate of AA binding to the resting enzyme (*V*_*31*_) has a high peak in an initial stage of the COX reaction, but its contribution to overall catalysis is negligible because of short duration of the peak (1 ms).

**Fig. 5.**
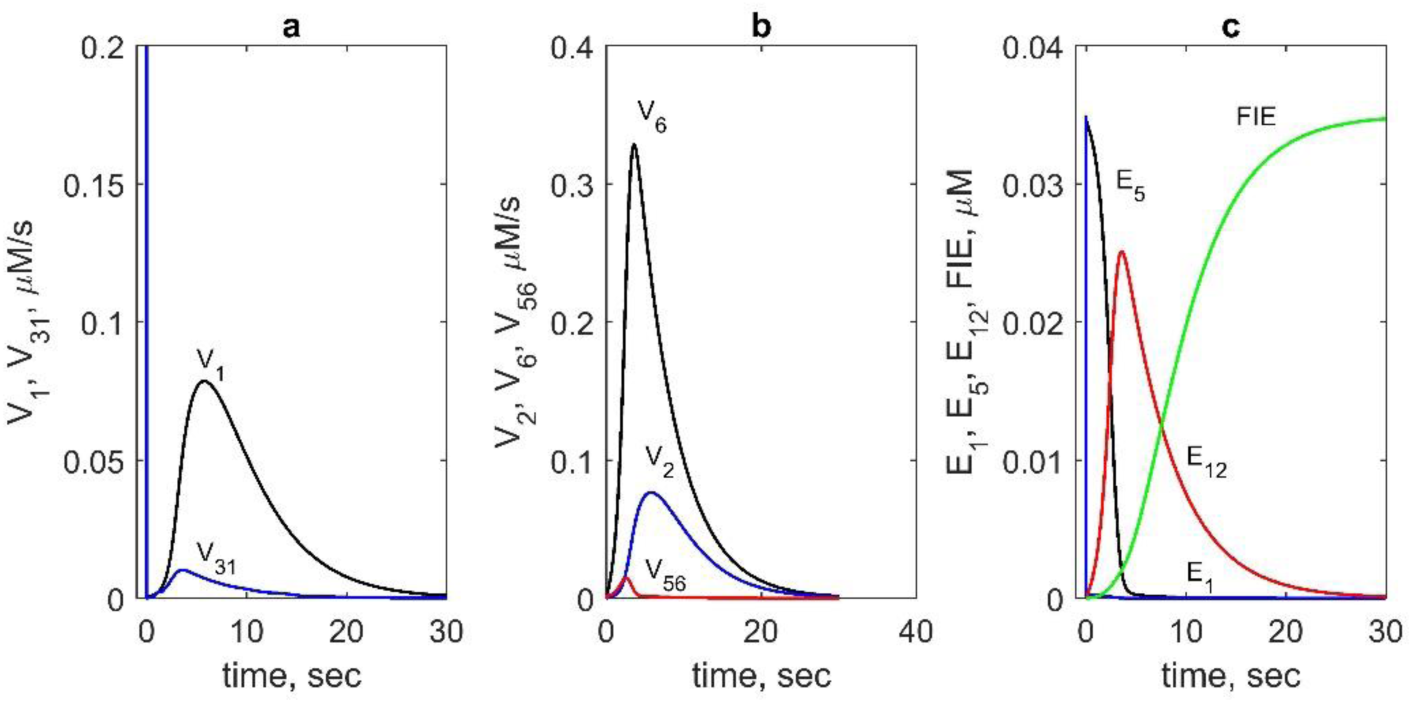
Contributions of different reactions and enzyme intermediates to PGHS-1 catalysis obtained in the model. (a) Reaction rates of AA binding to E_5_ state (V_1_, blue line) and the resting state E_1_ (V_31_, black line). (b) Reaction rates of AA radical formation: V_2_ (blue line), V_6_ (black line) and V_56_ (red line). (c) Kinetics of the main intermediates: E_1_ (blue line), E_5_ (black line), E_12_ (red line), and FIE (green line). In calculation: 35 nM PGHS-1, 80 μM AA and 100 μM RC.

The results of modelling also showed that the rate constant of electron transfer from Tyr385 residue to heme group depends significantly on the presence of AA in the COX site (reactions 13 and 18). The rate constant *k*_*9*_ of reaction 13 going between enzyme states with the empty COX site equals to 310 s^-1^, but the constant of reaction 18 between states with an occupied COX site *k*_*11*_ is much lower (1.1 s^-1^). Thus, reduction of heme group due to Tyr385 radical formation in the presence of AA in the COX site occurs weaker than that in the case of the empty COX site. Therefore, substrate-enzyme complex [Fe(IV), PP, Tyr*, AA] (E_9_) is formed mainly from Intermediate II according to the experimental data [5].

To determine what COX_i_ cycles contribute significantly to the overall COX reaction, we compared the reaction rates of the PGG_2_ synthesis in various COX_*i*_ cycles (Table 3) at the different concentration of RC (10 μM, 100 μM and 1000 μM). It turned out that the key COX cycle, in which maximum amount of PGG_2_ is produced, is the COX_2_ cycle (reaction 8). Recall that in our model heme group of the COX_2_ cycle is in the ground ferric state [Fe(III), PP], as opposite to the kinetic models of PGHS-1 [5], [9], where heme group is assumed to remain in the unreduced ferril state [Fe(IV), PP] throughout COX reaction (the COX_1_ cycle in our model).

**Table 3.** Reaction rates of PGH_2_ and PGG_2_ production (μM/s) in the different POX_i_ and COX_i_ cycles at the different concentrations of RC.

| RC, μM | $V_{11}$,<br>POX<sub>1</sub> | $V_{16}$,<br>POX<sub>2</sub> | $V_{23}$,<br>POX<sub>4</sub> | $V_4$,<br>COX<sub>1</sub> | $V_8$,<br>COX<sub>2</sub> | $V_{52}$,<br>COX<sub>3</sub> | Connection of COX<sub>1</sub> and<br>COX<sub>2</sub> cycles | | |
| --- | --- | --- | --- | --- | --- | --- | --- | --- | --- |
| | | | | | | | $V_{22}$ | $V_{28}$ | $V_{29}$ |
| 10 | 0.01 | 0.05 | 0.2 | 0.35 | 0.4 | $7 \cdot 10^{-3}$ | 3.35 | $10^{-3}$ | $10^{-3}$ |
| 100 | 0.01 | 0.05 | 0.3 | 0.07 | 0.63 | 0.01 | 6.94 | $5 \cdot 10^{-3}$ | $5 \cdot 10^{-3}$ |
| 1000 | 0.01 | 0.06 | 0.5 | 0.01 | 0.69 | 0.02 | 10.55 | 0.01 | 0.01 |

Calculation of the reaction rates showed that the COX_1_ cycle plays a significant role only at low concentrations of RC<10 μM (Table 3). This effect results from the increasing connection between COX_1_ and COX_2_ cycles that causes the reduction of heme group in the COX_1_ cycle by RC (reactions 19, 22, 28, and 29) and a decrease of the role of the COX_1_ cycle in the overall catalysis. Additionally, the COX_2_ cycle significantly contributes to the activation of AA oxygenation reaction in comparison with that in the COX_1_ cycle.

We also compared the rates of AA radical formation due to abstraction of a hydrogen atom from AA by Tyr385 radical in both COX cycles (Fig. 5b). The reaction rate *V*_*6*_ in the COX_2_ cycle was found to be significantly higher than the corresponding rate *V*_*2*_ in the COX_1_ cycle. We also compared contribution of two different reactions of AA radical formation which can happen either due to the Tyr385 radical according to the BC mechanism of PGHS-1 catalysis or due to another oxidant, oxyferryl heme, according to the early version of TC mechanism. The calculation showed that the reaction rate V_56_ of AA* formation owing to oxyferryl heme electron-transfer significantly lower that the corresponding reaction V_6_ proceeding due to Tyr385 radical (BC mechanism) (Fig. 5b).

To analyse the main reactions contributing considerably to PGH_2_ synthesis, we calculated the fluxes in the POX_i_ cycles, in which PGH_2_ is produced (Table 3). The main contribution to PGH_2_ production turned out to be made by reaction 23 in the POX_4_ cycle. Note that in the PGHS-1 models developed on a basis of the BC mechanism, PGH_2_ is produced in reaction 11 in the POX_1_ cycle. By contrast, in our model, reaction rate V_11_ is negligibly weak in comparison with reaction rate V_23_ (Table 3). To analyse this effect, we calculated the population kinetics of the enzyme states. Fig. 5c shows the time dependence of population of the states, maximum contributing to the PGHS-1 catalysis. These results showed that the POX_1_ cycle contributes noticeably at the initial stage of PGH_2_ synthesis, when the population of the resting state of the enzyme is maximum. In contrast, state E_12_ in the POX_4_ cycle works throughout all the catalysis, indicating that the POX_4_ cycle is active up to full SI of PGHS-1. Moreover, state E_12_ ([Fe(III), PP, Tyr, PGG*] being common for the COX_2_ and POX_4_ cycles participates simultaneously in the production of the intermediate product, PGG_2_ (reaction 23) as well as the final product PGH_2_ (reaction 8). As discussed above, these reactions make maximum contributions to the synthesis of PGG_2_ and PGH_2_ (Tables 3). Thus, in our model, COX_2_ and POX_4_ cycles are the main ones in PGHS-1 catalysis, while the COX_1_ and POX_1_ cycles contribute only at the initial stage of the PGHS-1 function.

### 2.5. Model application to the investigation of regulatory role of AA and RC in PGHS-1 function

The calibrated model was applied to study dependence of the PGHS-1 activity on substrate AA and cosubstrate RC concentrations. For comparison of the theoretical results with experiments we used validation experimental data which were not used in the fitting procedure. The comparison of model and experiment results made without any optional fitting was a validation test for the predictive power of the model. This allows the model to be applied to the reliable analysis of *in vivo* experimental data on the NSAD action.

First, we applied the model to calculate the dependence of the oxygen consumption rate on AA concentration, *V*_*O2*_(*AA*). For this, we first calculated kinetics of oxygen consumption rate, *V*_*O2*_(*t*), and compared the results with experimental data [8] (Fig. 6a). As seen, the model satisfactory describes experimental data on *V*_*O2*_(*t*) and reproduces the time of the enzyme activity, about 20 sec, that is in agreement with the SI time of PGHS-1 [1]. To plot the dependence of *V*_*O2*_(*AA*), we took the maximum values of the rates *V*_*O2*_(*t*) calculated at different AA concentrations. The theoretical dependence of *V*_*O2*_(*AA*) along with experimental data [11] are given in Fig. 6b.

**Fig. 6.**
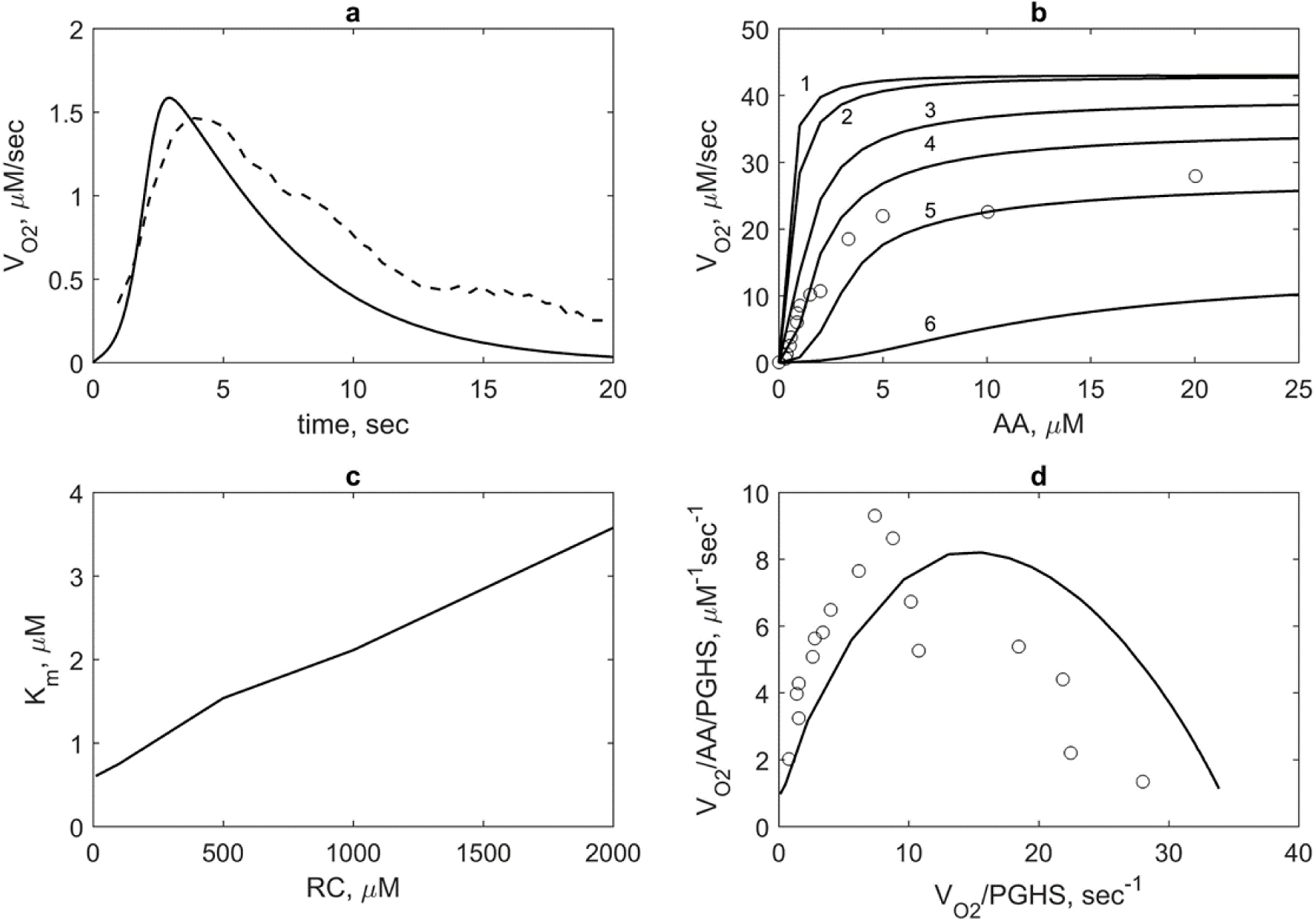
Model validation based on experimental data on oxygen consumption by PGHS-1. (a) Theoretical (solid line) and experimental (dash line) dependence of oxygen consumption rate V_O2_(t). Theoretical results obtained at 35 nM PGHS-1, 80 μM AA, and 300 μM RC; experimental data obtained at 35 nM PGHS-1, 80 μM AA and 100 μM ferulic acid [8]. (b) Dependence of oxygen consumption V_O2_ on AA concentration calculated at 10 μM (line 1), 100 μM (line 2), 500 μM (line 3), 1000 μM (line 4), 2000 μM (line 5), and 5000 μM (line 6) μM phenol; points – experimental data obtained at 1000 μM phenol and 0.005 μM PGHS-1 [11]. (c) Dependence of Michaelis constant K_m_ for AA substrate on RC concentration calculated in the conditions given in Fig. 6b. (d) The Eadie-Scatchard plot of value V_O2_/AA/PGHS on value V_O2_/PGHS. Lines and points – theoretical and experimental results obtained in the conditions provided in Fig. 6b.

We also calculated *V*_*O2*_(*AA*) at different RC concentrations (Fig. 6b). As seen, RC inhibits COX activity of the enzyme, i.e. an increase in RC concentration leads to an increase of apparent Michaelis constant *K*_*m*_ and decrease of the maximum oxygen consumption rate. This means that AA and RC compete each other for the COX site according to reactions 1 and 9 in the catalytic cycle (Fig. 2). Based on these results we calculated the dependence of the Michaelis constant *K*_*m*_ for AA on RC concentrations, *K*_*m*_(RC) (Fig. 6c). The linear dependence *K*_*m*_(RC) looks like the dependence of apparent Michaelis constant for substrate on competitive inhibitor concentration, where RC plays a role of an inhibitor. The range of theoretical values of *K*_*m*_ embraces a wide range of the experimental values of Michaelis constant, 3 μM [1], 5 μm [9] and 10.2 ± 1.5 μM [26]. Note that dissociation constants of AA binding with the enzyme obtained in our calculation is significantly lower, *K*_*d1,AA*_ equal to 0.1 μM than *K*_*m*_ (Table 2).

In the model, RC competes with AA for binding to states E_5_ and E_15_ in the COX_1_ cycle (reactions 1 and 9) and COX_2_ cycle (reactions 5 and 50), respectively. According with the BC mechanism, redaction of Tyr385 radical by RC leads to inhibition of COX activity.

Moreover, the discussed mechanism may explain the distinction of the obtained kinetic constants of the reduction rate of Tyr385 radical depending on a state of the COX site. In the parameter set (Table 2), the rate constant *k*_*10*_ of reactions 14 and 51 of Tyr385 reduction in states E_9_ (COX_1_ cycle) and E_20_ (COX_2_ cycle) with AA in the COX site equals approximately zero in contrast to the corresponding constant *k*_*5*_ in reaction 9 and 50 of Tyr385* reduction in states E_5_ (COX_1_ cycle) and E_15_ (COX_2_ cycle) with the empty COX site (Table 2). AA in the COX site is proposed not to permit RC to react with Tyr385*. For similar reaction 49 of Tyr385* reduction in states E_4_ (POX_2_-cycle) with PGG2* in the COX site, the constant equals *k*_*5*_, that may mean that RC can reduce two radicals state in the COX site.

An increase of *K*_*m*_ with increasing RC concentration is accompanied by a change in a curve shape of *V*_*O2*_(*AA*) from hyperbolic to the sigmoidal one at high RC concentrations (Fig. 6b). To investigate this positive cooperativity in PGHS-1 function, we plotted the Eadie-Scatchard dependence of ratio *V*_*O2*_/*AA/PGHS* on ratio *V*_*O2*_*/PGHS* (Fig. 6d) and provided the corresponding experimental data [11]. As seen, the theoretical downward-concave curve *V*_*O2*_/*AA/PGHS* vs. *V*_*O2*_*/PGHS* significantly differs from the linear Eadie-Scatchard plot corresponding to Michaelis kinetics (*V*_*O*2_*/AA =* − *V*_*o2*_*/K*_*m*_ *+ V*_*max*_*/K*_*m*_) that confirmed positive cooperativity in COX activity of PGHS-1 [26].

Positive cooperativity in the COX reactions defines the activation threshold of the PGHS-1 catalysis which depends on both AA and RC concentrations [11], [13]. Fig. 7 shows the activation threshold curve on a plan of AA and RC concentrations which divides the range of AA and RC concentrations on two regions, i.e. PGHS-1 is inactive and active at values of AA and RC concentrations lying below and upper the curve, respectively. Increasing activation threshold at increasing RC concentration is determined by inhibition of the autocatalytic COX reaction by RC. The PGHS-1 kinetics at AA and RC concentrations far from the activation threshold is close to classical Michaelis-Menten kinetics that is typical for *in vitro* experiments on PGHS-1 kinetics and experiments on its inhibition by NSAIDs. Nonlinear effect in PGHS-1 kinetics becomes more pronounced near the activation threshold, in the range of low AA concentrations (0.05 μM – 1 μM) which are likely to relate to cellular conditions [1]. Below this threshold, the PGHS-1 is in latent state and can be activated either increasing AA or decreasing RC concentrations as shown by arrows in Fig. 7.

**Fig. 7.**
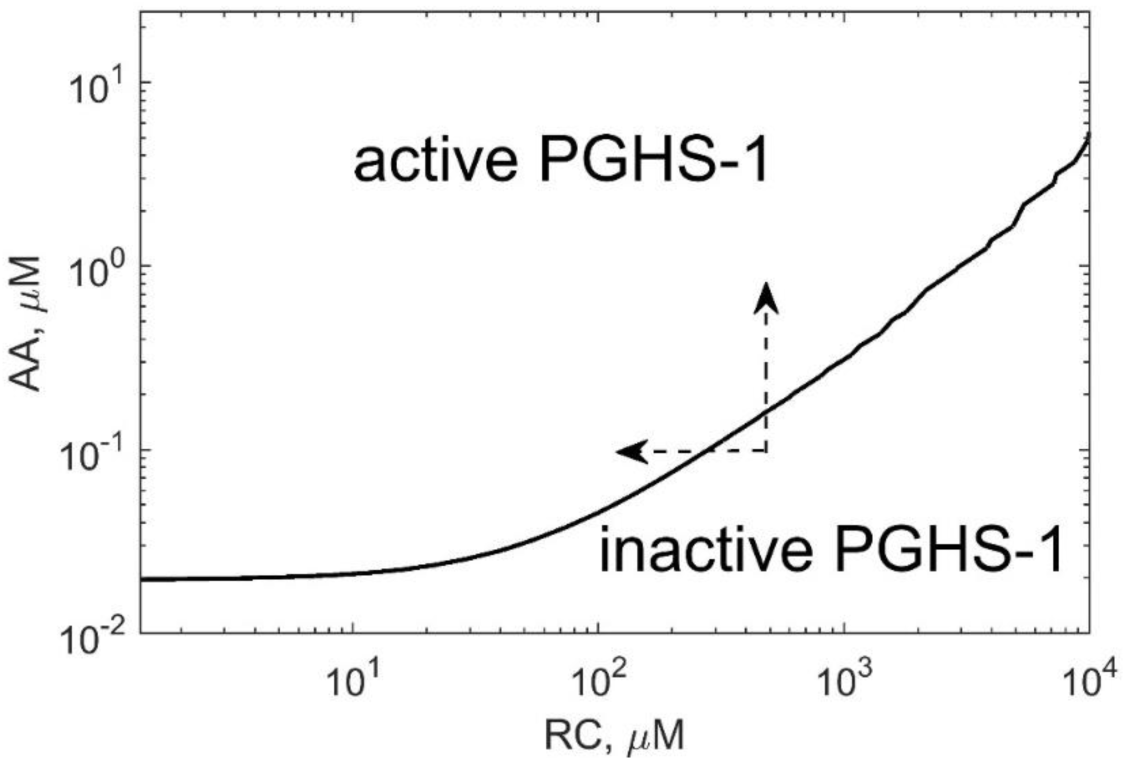
The activation threshold of PGHS-1 depending on AA and RC concentrations. The curve corresponds to PGH_2_ level of 0.02 μM.

### 2.6. Effect of reducing cosubstrate on COX activity of PGHS-1

We also applied our model to describe experimental data on the effect of RC on the oxygen consumption by PGHS-1 [10]. The results of calculation along with experimental data on the oxygen consumption rate, *V*_*O2*_, depending on phenol concentration are shown in Fig. 8a. As seen, RC practically does not affect the oxygen consumption rate at RC concentration less than 1000 μM and considerably inhibits COX activity at RC concentration higher than 1000 μM.

**Fig. 8.**
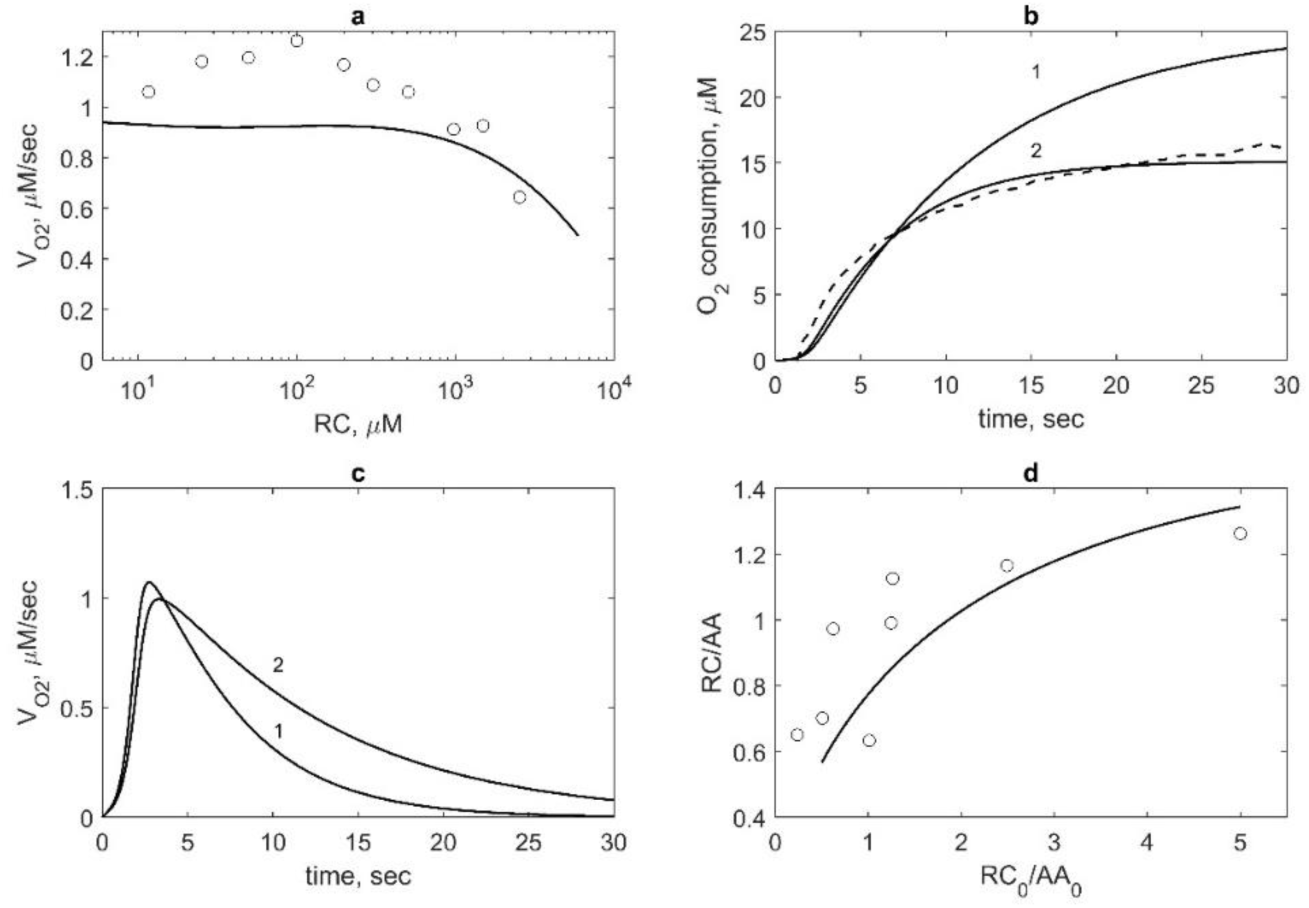
Model validation based on the experimental data on the RC (phenol) effect on PGHS-1 kinetics. (a) Theoretical (line) and experimental (points) [10] dependence of oxygen consumption rate V_O2_ on RC concentration. V_O2_ is normalised by its value obtained in the absence of RC. Calculation and experiment were carried out at 47 nM PGHS-1 and 100 μM AA. (b) Theoretical (solid lines) and experimental (dash line) kinetics of oxygen consumption. Calculation was carried out at 47 nM PGHS-1, 100 μM AA, 1000 μM RC (line 1) and 200 μM RC (line 2). Experimental data were obtained at 47 nM PGHS-1, 100 μM AA, and 1000 μM phenol [10]. (c) Dependence of oxygen consumption rate V_O2_ calculated at 47 nM PGHS-1, 100 μM AA, 1000 μM RC (line 1) and 200 μM RC (line 2). (d) Dependence of the ratio of RC and AA consumption, RC/AA, on the ratio of initial concentrations, RC_0_/AA_0_. Line – calculation at 35 nM PGHS-1 and points – experimental data obtained 35 nM PGHS-1, 20 μM, 40 μM, and 80 μM AA and 20 μM, 50 μM, and 100 μM ferulate [8]. The ratio RC/AA was taken at 30 s after reaction initiation.

For the comparison of these theoretical results with experimental data we used experimental data corrected for oxygen electrode signal [10]. Note that these data show a slight increase of *V*_*O2*_, approximately at 20 % above the control values in the comparison with uncorrected data [10] and data obtained by another group [24] which have the pick of *V*_*O2*_ about 2-3 times the control value. Our result showing no stimulation effect of RC on the COX reaction are closer to the corrected experimental and theoretical data [10].

According to our model, RC has an indirect stimulation effect on oxygen consumption. Calculation of O_2_ consumption kinetics showed its increase at growing RC concentration (Fig. 8b). In the model the enhancement of O_2_ consumption at RC concentration up to 1000 μM results from the extension of the functioning time of the enzyme due to its protecting from SI by RC. Increasing of the functioning time of PGHS-1 is seen well in Fig. 8c where time dependence of *V*_*O2*_ at different RC concentrations is shown.

### 2.7. Analysis of the connection between COX and POX activities

To analyse how our model describe stoichiometry between cosubstrate oxidation and arachidonic acid oxygenation during PGHS-1 functioning, we calculated dependence of the ratio between the rate of cosubstrate (phenol) and arachidonate consumption, RC/AA, on the ratio between their initial concentrations, RC_0_/AA_0_. Comparison of the theoretical results with experimental data [8] are shown in Fig. 8d.

The dependence of RC/AA shows the extent of the interconnection between the POX and COX cycles in the course of PGHS-1 function. The obtained ratio RC/AA is ranged from 0.5 to 1.3 that is consistent with the experimental range (0.6-1.3) [8], when RC/AA is in the region of 0.5 - 5. The theoretical data agree with the prediction of the BC mechanism of PGHS-1 catalysis (0-2) [8], [19]. In our calculation, the dependence of RC/AA on RC_0_/AA_0_ is the dependence with saturation rather than linear dependence [8].

As AA consumption defines PGG_2_ production, and RC consumption relates to PGH_2_ synthesis, the ratio RC/AA determines the extent of the PGG_2_ conversion to PGH_2_. Because complete conversion of PGG_2_ to PGH_2_ requires 2 molecules of RC (RC/AA=2), the obtained value of the ratio RC/AA= 0.5-1.3 shows that POX cycles are slower than the COX ones and complete conversion of PGG_2_ is not reached. Thus, according to the model, some fraction of the intermediate product, PGG_2_, remains in solution at the end of the catalytic cycle and is accumulated there. Note, the accumulation of PGG_2_ in solution is proposed to be important for activation of PGHS-1 at cell stimulation.

## 3. Discussion

The microscopic network model of PGHS-1 catalysis was developed as the further extension of a series of the models developed previously on a basis of the Branch Chain mechanism of PGHS-1 kinetics [8], [9], [10]. Our model presents a network of the interconnected five POX and three COX cycles proceeding independently of each other and including all possible microscopic states generated in the POX and COX sites and possible transitions among these multiple states during catalysis. Analysis of the cycle structure of this network model of PGHS-1 was carried out by Yordanov and Stelling [27] on a basis of the developed graph-based approach to the investigation of combinatorial complexity of enzymatic kinetics and signalling network dynamics.

Model calibration was carried out an a basis of the experimental datasets obtained at the homogeneous experimental conditions of PGHS-1 treated by AA and phenol as RC [11] that allowed consistent description of a majority of kinetic data known for PGHS-1 catalysis under wide experimental conditions within a unified set of kinetic parameters. The calibrated model reproduced satisfactorily key features of the complex PGHS-1 dynamics, i.e. interaction of COX and POX activities, processes of enzyme self-inactivation, its autocatalytic mechanism, and the activation threshold of the enzyme. Correct description of these characteristics of PGHS-1 in the framework of self-consistent kinetic model with the unified set of kinetic parameters is critical for *in silico* screening and prediction of the NSAID inhibition effects in different experimental assays and targeted cells.

The developed model was also applied to the experimental conditions, where PGHS-1 was treated by other substrates, adrenaline as RC and H_2_O_2_ as a substate of the POX site [12] As a result, a new subset of the parameters responsible for binding of these substrates to the enzyme was identified (Table 2). This showed that for the further application of the model to experimental conditions different from those used in this work (e.g. other cosubstrates), it is necessary to identify a subset of the kinetic parameters related to the binding of other substrates to PGHS-1 and fix the rest of the kinetics parameters characterising intra-enzymatic kinetics obtained in the fitting procedure (Table 2). Moreover, identification of the kinetic parameters of hydrogen peroxide reaction with PGHS-1 in the model (Table 2) allows the mode to be applied to the investigation of the PGHS-1 kinetics and NSAID effectiveness at different hydrogen peroxide levels in various cells and organs [15].

To validate and test predictive power of the developed model we applied it to the description of regulatory properties of the PGHS-1 and compared the results with additional experimental data not used in the calibration procedure. A study of the enzyme activity showed that AA and RC play various regulatory roles. We elucidated different mechanisms of inhibition and activation effects of RC on the enzyme. AA along with RC was shown to play activation role, switching the enzyme from an inactive state to the functioning one. In this case, RC was found to influence the activation threshold of the enzyme. Note that detailed understanding and description of these regulatory properties are important for the correct analysis of platelets activation as well as NSAID effects on this process.

The developed model was applied to study intermediate enzyme states and reactions which contribute significantly to the synthesis of PGG_2_ and PGH_2_. Taking into account two basic mechanisms of PGHS-1 function in the model, i.e. the Branched Chain [5], [8] and Tightly Coupled [6], [7] mechanisms allowed us to estimate the contribution of each mechanism to the prostaglandin H2 biosynthesis.

The model supported the BC mechanism of PGHS-1 function, which assumes that Tyr385 radical in the COX site is a key component in interconnection between the COX and POX activities [8]. We also concluded that Tyr385 radical formation is required for effective binding of AA to the COX site. For AA to bind effectively, PGHS-1 first should be activated by a formation of Tyr385 radical due to POX activity of the enzyme. This effect is likely to result from a possible conformation change in the COX site by Tyr385 radical formation that can lead to an increase of the COX site affinity to AA. Conversely, AA in the COX site can suppress Tyr385 radical formation. Note that the experimental data indicates the possible mutual interaction between molecular compound in the COX site and Tyr385 radical formation. For example, experimental studies strongly indicate that the tyrosyl radical in inhibitor-treated PGHS-1 is probably not located at Tyr385 [28], and the alternative tyrosyl radical is not competent for COX catalysis. This data can serve an example of the effect of inhibitor molecules on the enzyme structure and internal dynamics particularly on the suppression of Tyr385 radical formation in the COX site [29].

As the result of the model fitting against the experimental data, dissociation constants *K*_*d,1*_ of AA binding to the states containing Tyr385 radical (E_5_ and E_15_ in the COX_1_ and COX_2_ cycles, respectively) was obtained to be lower than the corresponding constants *K*_*d,12*_ of AA binding to the states not containing Tyr385 radical (E_1_, E_2_ and E_3_ states in the POX_1_ cycle, see Table 2). A negligible contribution of reactions 30-32 to the catalysis did not permit studying carefully the difference in dissociation constants of AA binding to various enzyme states. The additional experimental and theoretical investigations are needed for elucidation of the Tyr385 radical influence on enzyme affinity to AA.

In the model, we also investigated different mechanisms of PGHS-1 irreversible self-inactivation (SI) which was suggested to be additional regulatory mechanism preventing excess generation of signalling molecules by the enzyme in cells [3]. The SI mechanism in the model included separate inactivation of the COX and POX activities resulting from the enzyme oxidative damage caused by radical intermediates generated in the POX and COX sites of PGHS-1 [20]. The model with the obtained reaction rates of SI satisfactorily described stopping kinetics of PGH_2_ production and full inactivation of the enzyme occurring within approximately 20 min (Figs. 3a, 3b and 6a).

The flux analysis of the network model showed the basic role of the COX_2_ cycle which includes the ferric state of the enzyme [Fe(III), PP] (E_15_). This conclusion contradicts the models where only ferril state [Fe(IV), PP] (E_5_) was considered in the COX reactions [5], [8]. To remove this contradiction, the further investigation of the connectivity between the COX_1_ and COX_2_ cycles is needed.

As it has been noticed in section 2.1, we considered in the model an alternative mechanism of the COX reaction initiation which was proposed in the framework of the early version of the TC mechanism [6]. According to this mechanism, a direct hydrogen atom abstraction from AA by the ferryloxo-protoporphyrin radical cation of the heme group (reaction 56 from the E_14_ state in POX_4_ cycle) occurs, when AA is in the COX site. The flux analysis allowed us to compare this mechanism of AA radical formation with the alternative mechanism through Tyr385 radical, proposed in framework of the BC mechanism [5]. The obtained values of maximum reaction rates *V*_*2*_ in the COX_1_ cycle, *V*_*6*_ in the COX_1_ cycle, *V*_*64*_ from the E_19_ state, and *V*_*56*_ showed that the main mechanism of AA radical formation is the BC mechanism. Thereby, the first stage of AA oxygenation including AA radical formation occurs mainly due to Tyr385* reduction in accordance with the BC mechanism.

To investigate the inhibition role of RC we calculated the dependence of the apparent Michaelis constant *K*_*m*_(RC) for AA on RC concentration and showed an increase of *K*_*m*_ with increasing RC concentration. The obtained variation of *K*_*m*_ can explain a wide range of its variation (3 μM - 5 μM) obtained in different experiments [1]. It is to be noted that there are not any reliable data on RC nature and its concentration in different cells and *in vitro* measurements of the dose dependence for NSAIDs are usually carried out at high RC concentration (e.g. 1000 μM of phenol). A change in the apparent Michaelis constant *K*_*m*_ for AA under RC concentration variation affects the values of IC_50_ for drugs measured at different RC and peroxide concentrations [16]. We also obtained dissociation constant *K*_*d,1*_ of AA (0.1 μM) which is an important value to estimation NSAIDs and AA competition for the COX site and correct analysis of drugs IC_50_s in various experimental conditions [15].

The inhibition effect of RC on PGHS-1 catalysis observed in the model at high concentration of RC (>1000 μM, Fig. 8a) can be explained by RC competition for Tyr385 radical in the COX site. RC molecule (e.g. phenol) is likely to be able to penetrate the COX site and reduce Tyr385 radical that abrogates oxygenation reaction. Note that the same mechanism was proposed for the interaction of nitric oxide with PGHS-1, i.e. **•**NO can terminate the COX activity by its reaction with Tyr385 radical forming 3-nitrotyrosine [30]. Similar mechanism is suggested to explain an inhibition effect of phenolic antioxidant, acetaminophen acting as reducing cosubstrate for the peroxidase protoporphyrin and preventing tyrosine radical formation in the COX site [31].

According to experimental data, RC also causes a stimulation effect on PGHS-1 catalysis at low RC concentration [24], [31]. Our model partly explained the stimulation mechanism of RC (phenol, acetaminophen and others) by preventing the enzyme from suicide inactivation and extending the catalytic time of PGHS-1 (Figs. 3a, 3b and 8b). Thus, our model is recommended to use in analysis of the PGHS-1 kinetics and NSAID effects in the range of high concentration of RC (>200 μM phenol). The further extension of the current model is required to study the stimulation role of RC in PGHS-1 catalysis at low concentration of RC.

Analysis of the oxygen consumption dependence on substrate and co-substrate concentrations showed its sigmoidal shape in the range of low AA and high RC concentrations (Fig. 6b) that points at positive cooperativity and nonlinear mechanism in PGHS-1 functioning. Positive cooperativity in the enzyme kinetics is also confirmed by the calculation of the Eadie-Scatchard plot that significantly differs from linear dependence and has downward-concave shape (Fig. 6d). The molecular mechanic of positive cooperativity of PGHS-1 function is not entirely understood. In the case of dimer structure of PGHS-1, a possible mechanism of cooperativity can be determined by either allosteric interaction between two subunits of the holoenzyme or cooperativity within each monomer [24], [26]. In our model, we proposed independent function of the PGHS-1 monomers, thus manifestation of cooperativity in the model does not connected with its dimer structure. An increase in the extent of cooperativity with increasing RC and decreasing AA concentrations suggested that the POX activity and the reducing co-substrate concentration are determinant factors in positive cooperativity in the COX reactions. Cooperativity of the COX activity in the model is linked to the autocatalytic effect of AA, because it is observed profoundly close to the activation threshold of PGHS-1. The mechanism of cooperativity is related to positive feedback via the POX cycle which is activated at high RC concentrations. Note that the positive cooperativity and noticeable activation threshold are absent in the inducible PGHS-2 isoform whose kinetics is closer to the classical Michaelis-Menten one [13], [26]. This difference in the initiation kinetics of two isoforms of PGHSs in the range of cellular submiromolar concentration of AA suggests a different control mechanism in prostaglandin biosynthesis by constitutive PGHS-1 and inducible PGHS-2. In case of constitutive PGHS-1, the activation threshold adjustable to cellular conditions ensures a fast activation of latent enzymes and their inactivation by triggering of AA concentration to cellular demands nearby the activation threshold. The lower activation threshold and the absence of positive cooperativity in initiation kinetic of PGHS-2 guarantees a fast and strong response of the enzyme to the cellular stimulus [13]. It is proposed that distinctive regulatory properties of two isoforms of PGHSs reflect their different physiological roles in the same cells expressing both PGHS-1 and PGHS-2 and consume the same substrate and cosubstrate.

## 4. Methods

### 4.1. Parametrisation of the PGHS-1 model

In accordance with the developed scheme for the PGHS-1 catalytic cycle (Fig. 2) we developed a mathematical model for PGHS-1 catalysis. The model includes 28 ordinary differential equations (ODEs) formulated for 24 enzyme catalytic states (E_i_) and six metabolite concentrations (AA, O_2_, RC, OC, PGG_2_, and PGH_2_). The whole system of ODEs and conservation laws are presented in Supplementary Materials. To run numerical simulation and calibration of the model we used the software package for kinetic modelling DBsolve 6.1 [32] and the Systems Biology Toolbox for MATLAB (Mathworks).

The reaction rate expressions for all the reactions were formulated according to the mass action law with the rate constants being the parameters of the model (see Supplementary Materials). To identify the model parameters, we fitted the solutions of the ODEs against the experimental data on enzyme kinetics of PGHS-1 available from the literature.

The model includes 66 reactions and 68 corresponding reaction rate constants. The experimental data published in literature are not sufficient for reliable identification of all the model parameters. The lack of the data is mainly due to some peculiarities of the experimental design generally used for kinetic studies of PGHS-1 catalysis. The majority of such experimental studies included either the time courses for metabolite concentrations or reaction rates of their consumption or production such as AA or O_2_ consumption, RC oxidation, and PGH_2_ production. Whereas these characteristics are relatively easy to study experimentally, the measurement of kinetic dependencies for individual enzyme intermediates (catalytic states E_i_ in our model) requires application of different experimental techniques, mainly EPR spectrometry for the radical intermediates detection [14]. To reliably identify 68 reaction rate constants, we would ideally need time series data for the majority of enzyme catalytic states considered in the model. Taking into account lack of such the data we proposed the following way to reduce the number of parameters in the model.

In general, all the reactions in the developed catalytic cycle shown in Fig. 2 can be classified into several groups, as some of them are of similar types. For example, we grouped together the reactions of AA binding to the COX site (reactions 1, 30, 31, and others), or reduction of PGG_2_ to PGH_2_ (reactions 11, 16, 20, and others). The reactions within the groups differ from each other by the enzyme catalytic state participating in corresponding reactions. For example, AA can bind to the enzyme being in the resting state [Fe(III), PP, Tyr] (E_1_, reaction 31) and ferryl states [Fe(IV), PP, Tyr] and [Fe(IV), PP*+, Tyr] (E_3_ and E_2_ states in reactions 30 and 32, respectively). Alternatively, AA can bind to the enzyme state containing Tyr385 radical (E_5_, E_15_ and E_11_ states in reactions 1, 5 and 47, respectively). Therefore, it was assumed that the reactions of similar type would have similar rate constants.

However, in general case kinetic constants of the elementary reactions taking place in one of the catalytic sites can depend on the state of all other catalytic components, located in the nearest proximity and involved in the further catalytic steps. For example, there are experimental evidences that AA binding to the COX site may depend on the presence/absence of the radical on Tyr385 [18]. PGHS-1 is a bifunctional enzyme and though the reactions traditionally referred to as COX activity and POX activity are catalysed in different sites, there are indications that the processes in one site can influence the rate of the processes in the other one. This phenomenon is traditionally discussed in the context of cooperativity of PGHS-1 [10].

We tried different approaches in grouping the model parameters and analysed the results of fitting of the corresponding models to the same set of experimental data. Analysis of the fitting results allowed us to hypothesise which reaction rates do not depend much on the processes taking place in the neighbour site, and therefore to conclude, which kinetic constants can be assumed to be equal to each other. This analysis also allowed us to make conclusions on the cooperativity between COX and POX activities. The approach we used finally allowed significantly reducing the number of parameters subject to fitting. The best fitting of the model to experimental data was achieved with the set of 18 parameters, given in Table 2.

When identifying the model parameters, we were pursuing the goal to find a unified set of parameters, which would allow consistently describing a majority of *in vitro* kinetic data known for PGHS-1 catalysis. Identification of the model parameters and their further validation was performed as follows. We have chosen several experimental data sets obtained in different research laboratories to fit the model parameters against the data, and then validated the calibrated model by checking its applicability to the description of experimental data which were not taken in the fitting procedure.

To avoid ambiguity, for the initial stage of the fitting, among all kinetic information available for PGHS-1, we have chosen three homogeneous datasets which were obtained for the same object, purified PGHS-1 from sheep seminal vesicles, under approximately the same experimental conditions (temperature and pH). The first set included kinetic data on PGG_2_ and PGH_2_ production (points in Figs. 3a and 3b) measured at the different concentration of RC, phenol [8]. As the second dataset, we used the kinetic data on AA consumption (points in Fig. 3c), obtained for the different AA concentrations at the fixed concentration of RC, phenol [11]. The experimental conditions of the third dataset [12] differed from the first two ones. In this experiment, kinetics of the separate POX activities of PGHS-1 treated by H_2_O_2_ as the substrate of the POX site and adrenaline as RC was measured in the absence of substrate AA. These data were used to identify kinetic parameters *k*_*5*_, *k*_*6*_ and *k*_*8*_ of the POX reactions for PGHS-1 treated by H_2_O_2_. The obtained values of these parameters were different from those identified from the experimental dataset on the kinetics of PGHS-1 treated by phenol as RC and AA (see Table 2). Note that calibration of the model based on two different experimental conditions allows the model to be applied to the analysis of experimental data obtained for PGHS-1 treated by the different RCs and different substrates of the POX site (e.g. PGG_2_ and H_2_O_2_).

### 4.2. Global sensitivity analysis of the model

Global sensitivity analysis (GSA) of the model to the kinetic parameters obtained in the fitting procedure was based on Morris GSA method of the calculation of the elementary effects [25], [33]. The resulting Morris ranking is calculated as

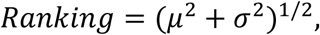

where μ and σ are two sensitivity measures in Morris GSA method, i.e. μ assesses the impact of a parameter on the model output and σ describes the non-linear effects and interactions of the parameters. As the model output in the calculation, we took an area under the curve (AUC) of PGH_2_ time course.

## 5. Conclusion

The key aim of this work was the development of reliable and validated kinetic model of PGHS-1 which reproduces main features of its complex kinetics and can be used as a computational tool for accurate prediction of inhibition effect of NSAIDs and their combinations in a range of cellular conditions. The cycle based network model integrates both the current models of the molecular mechanism of PGHS-1 catalysis and a large dataset of experimental data on the kinetics and regulatory properties of PGHS-1. This integrative approach helped us to overcome the limitation of the previously developed PGHS-1 models mainly applicable to the description of individual *in vitro* data in a specific range of substrate and cosubstarte concentrations.

The developed model of PGHS-1 catalysis is a platform for the further development of a kinetic model of inducible isoform PGHS-2 (COX-2) which is expressed together with PGHS-1 in vascular endothelia cells and other organs and is a target of COX-2 selective NSAIDs. Development of the kinetics models of two isoforms within an unified integrative approach would allow investigation of the different physiological roles of these isoforms in the same cells and analysis of NSAID inhibition effect on the balance of pro-thrombotic (thromboxane) and anti-thrombotic (prostacyclin) prostaglandins in platelets and endothelial cells.

## Supplementary Materials

Supplementary material contains the ODEs of the PGHS-1 model.

## Acknowledgement

Authors acknowledge Dr Galina Lebedeva and Dr Oleg Demin for fruitful discussion and suggestions on the modelling strategy.

## Author Contributions

Conceptualization, AG and YK; Methodology, AG and YK; Software, AG and MS; Validation, AG; Formal Analysis, KP; Investigation, AG, YK, MS, and KP; Data Curation, KP; Writing – Original Draft Preparation, AG; Writing – Review & Editing, AG, YK, MS, and KP; Visualization, MS; Supervision, YK; Project Administration, KP.

