## Supplemental Materials for "Cycle network model of Prostaglandin H Synthase-1"

### Supplementary Materials

#### Ordinary differential equations of the PGHS-1 model

$$\frac{dE_1}{dt} = V_{10} - V_{11} - V_{31} + V_{50} - V_{72} \tag{1}$$

$$\frac{dE_2}{dt} = V_{11} - V_{12} - V_{13} - V_{32} - V_{38} \tag{2}$$

$$\frac{dE_3}{dt} = V_9 - V_{10} + V_{12} - V_{30} \tag{3}$$

$$\frac{dE_4}{dt} = V_{35} - V_{37} - V_{49} + V_{54} - V_{56} \tag{4}$$

$$\frac{dE_5}{dt} = -V_1 + V_4 - V_9 + V_{13} - V_{28} - V_{43} + V_{53} + V_{57} - V_{58} \tag{5}$$

$$\frac{dE_6}{dt} = V_{14} - V_{15} + V_{17} + V_{30} \tag{6}$$

$$\frac{dE_7}{dt} = V_{23} - V_{24} + V_{33} - V_{35} - V_{41} - V_{52} \tag{7}$$

$$\frac{dE_8}{dt} = V_3 - V_4 - V_{22} + V_{24} + V_{49} \tag{8}$$

$$\frac{dE_9}{dt} = V_1 - V_2 - V_{14} + V_{18} - V_{29} - V_{44} - V_{61} \tag{9}$$

$$\frac{dE_{10}}{dt} = V_{15} - V_{16} + V_{31} + V_{50} \tag{10}$$

$$\frac{dE_{11}}{dt} = -V_{47} + V_{52} - V_{53} - V_{54} \tag{11}$$

$$\frac{dE_{12}}{dt} = V_7 - V_8 + V_{22} - V_{23} \tag{12}$$

$$\frac{dE_{13}}{dt} = V_2 - V_3 - V_{19} + V_{21} + V_{48} + V_{56} \tag{13}$$

$$\frac{dE_{14}}{dt} = V_{16} - V_{17} - V_{18} + V_{32} - V_{39} - V_{56} \tag{14}$$

$$\frac{dE_{15}}{dt} = -V_5 + V_8 + V_{28} - V_{45} - V_{50} - V_{60} \tag{15}$$

$$\frac{dE_{16}}{dt} = V_6 - V_7 + V_{19} - V_{20} \tag{16}$$

$$\frac{dE_{17}}{dt} = V_{34} - V_{36} - V_{48} - V_{54} \tag{17}$$

$$\frac{dE_{18}}{dt} = V_{25} - V_{26} + V_{60} + V_{59} \tag{18}$$

$$\frac{dE_{19}}{dt} = V_{47} - V_{64} - V_{65} - V_{66} - V_{67} - V_{68} \tag{19}$$

$$\frac{dE_{20}}{dt} = V_5 - V_6 + V_{29} - V_{46} - V_{51} - V_{59} \tag{20}$$

$$\frac{dE_{21}}{dt} = V_{20} - V_{21} - V_{33} - V_{34} - V_{40} \tag{21}$$

$$\frac{dE_{22}}{dt} = -V_{25} + V_{27} + V_{36} + V_{37} + V_{58} + V_{61} \tag{22}$$

$$\frac{dE_{23}}{dt} = V_{26} - V_{27} - V_{42} \tag{23}$$

$$\frac{dFIE}{dt} = V_{38} + V_{39} + V_{40} + V_{41} + V_{42} + V_{43} + V_{44} + V_{45} + V_{46} + V_{54} \tag{24}$$

$$\frac{dAA}{dt} = -V_1 - V_5 - V_{30} - V_{31} - V_{32} \tag{25}$$

$$\frac{dO_2}{dt} = -2 \cdot V_3 - 2 \cdot V_7 - 2 \cdot V_{33} - 2 \cdot V_{55} \tag{26}$$

$$\frac{dPGH_2}{dt} = V_{11} + V_{16} + V_{20} + V_{23} + V_{26} \tag{27}$$

$$\frac{dRC}{dt} = -V_9 - V_{10} - V_{12} - V_{14} - V_{15} - V_{17} - V_{19} - V_{21} - V_{22} - V_{24} - V_{25} - V_{27} - V_{28} - V_{29} - V_{48} - V_{49} - V_{50} - V_{51} - V_{53} \quad (28)$$

$$\sum_{i=1}^{24} E_i = E_o \quad (29)$$

$$RC + OC = RC_O \quad (30)$$

$$PGH_2 + PGG_2 = AA_0, \quad (31)$$

where rate equations $V_i$ are defined by the following relations:

$V_1 = k_1 \cdot (E_5 \cdot AA - K_1 \cdot E_9)$ (32)
$V_2 = k_2 \cdot E_9$ (33)
$V_3 = k_3 \cdot E_{13} \cdot O_2 \cdot O_2$ (34)
$V_4 = k_4 \cdot E_8$ (35)
$V_5 = k_1 \cdot (E_{15} \cdot AA - K_1 \cdot E_{20})$ (36)
$V_6 = k_2 \cdot E_{20}$ (37)
$V_7 = k_3 \cdot E_{16} \cdot O_2 \cdot O_2$ (38)
$V_8 = k_4 \cdot E_{12}$ (39)
$V_9 = k_5 \cdot E_5 \cdot RC$ (40)
$V_{10} = k_6 \cdot E_3 \cdot RC$ (41)
$V_{11} = k_7 \cdot PGG_2 \cdot E_1$ (42)
$V_{12} = k_8 \cdot E_2 \cdot RC$ (43)
$V_{13} = k_9 \cdot E_2$ (44)
$V_{14} = k_{10} \cdot E_9 \cdot RC$ (45)
$V_{15} = k_6 \cdot E_6 \cdot RC$ (46)
$V_{16} = k_7 \cdot PGG_2 \cdot E_1$ (47)
$V_{17} = k_8 \cdot E_{14} \cdot RC$ (48)
$V_{18} = k_{11} \cdot E_{14}$ (49)
$V_{19} = k_6 \cdot E_{13} \cdot RC$ (50)
$V_{20} = k_7 \cdot PGG_2 \cdot E_{16}$ (51)
$V_{21} = k_8 \cdot E_{21} \cdot RC$ (52)
$V_{22} = k_6 \cdot E_8 \cdot RC$ (53)
$V_{23} = k_7 \cdot PGG_2 \cdot E_{12}$ (54)
$V_{24} = k_8 \cdot E_7 \cdot RC$ (55)
$V_{25} = k_6 \cdot E_{22} \cdot RC$ (56)
$V_{26} = k_7 \cdot PGG_2 \cdot E_{18}$ (57)
$V_{27} = k_8 \cdot E_{23} \cdot RC$ (58)
$V_{28} = k_6 \cdot E_5 \cdot RC$ (59)
$V_{29} = k_6 \cdot E_9 \cdot RC$ (60)
$V_{30} = k_{12} \cdot (E_3 \cdot AA - K_{12} \cdot E_6)$ (61)
$V_{31} = k_{12} \cdot (E_1 \cdot AA - K_{12} \cdot E_{10})$ (62)
$V_{32} = k_{12} \cdot (E_2 \cdot AA - K_{12} \cdot E_{14})$ (63)
$V_{33} = k_3 \cdot E_{21} \cdot O_2 \cdot O_2$ (64)
$V_{34} = k_{13} \cdot E_{21}$ (65)
$V_{35} = k_{14} \cdot E_7$ (66)
$V_{36} = k_{15} \cdot E_{17}$ (67)
$V_{37} = k_{16} \cdot E_4$ (68)
$V_{38} = k_{in1} \cdot E_2$ (69)
$V_{39} = k_{in1} \cdot E_{14}$ (70)
$V_{40} = k_{in1} \cdot E_{21}$ (71)
$V_{41} = k_{in1} \cdot E_7$ (72)
$V_{42} = k_{in1} \cdot E_{23}$ (73)
$V_{43} = k_{in2} \cdot E_5$ (74)
$V_{44} = k_{in2} \cdot E_9$ (75)
$V_{45} = k_{in2} \cdot E_{15}$ (76)
$V_{46} = k_{in2} \cdot E_{20}$ (77)
$V_{47} = k_1 \cdot (AA \cdot E_{11} - K_1 \cdot E_{19})$ (78)
$V_{48} = k_5 \cdot E_{17} \cdot RC$ (79)
$V_{49} = k_5 \cdot E_4 \cdot RC$ (80)
$V_{50} = k_5 \cdot E_{15} \cdot RC$ (81)
$V_{51} = k_{10} \cdot E_{20} \cdot RC$ (82)
$V_{52} = k_4 \cdot E_7$ (83)
$V_{53} = k_8 \cdot E_{11} \cdot RC$ (84)
$V_{54} = k_{in1} \cdot E_{11}$ (85)
$V_{55} = k_3 \cdot E_{17} \cdot O_2 \cdot O_2$ (86)
$V_{56} = k_{13} \cdot E_{14}$ (87)
$V_{57} = k_{14} \cdot E_4$ (89)
$V_{58} = k_{in} \cdot E_5$ (90)
$V_{59} = k_{in} \cdot E_{20}$ (91)
$V_{60} = k_{in} \cdot E_{15}$ (92)
$V_{61} = k_{in} \cdot E_9$ (93)
$V_{62} = k_{in} \cdot E_{11}$ (94)
$V_{63} = k_7 \cdot PGG_2 \cdot E_{15}$ (95)
$V_{64} = k_2 \cdot E_{19}$ (96)
$V_{65} = k_5 \cdot E_{19} \cdot RC$ (97)
$V_{66} = k_8 \cdot E_{19} \cdot RC$ (98)
